# Lateral root formation is stimulated by common symbiosis genes and *NIN* in *Lotus japonicus*

**DOI:** 10.64898/2025.12.16.694690

**Authors:** Chloé Cathebras, Aline Sandré, Martina K. Ried-Lasi, Martin Parniske

## Abstract

Lateral roots (LR) and the root nodules (RN) of legumes are structurally related and the decision processes leading to RN formation involve signal exchange with the shoot. In order to disentangle these processes, we established a quantitative assay for LR formation in hairy root liquid cultures (HRLC) for the legume *Lotus japonicus*. In HRLC, ectopic expression of *SymRK*, or deregulated, auto-active versions of CCaMK and Cyclops stimulated LR formation in a *NIN*-dependent manner, but spontaneous RN were never observed. It appears that the previously described spontaneous RN formation induced by these versions requires the presence of the shoot. Interestingly, CCaMK^T265D^ increased LR number in a *cyclops* mutant, revealing the presence of additional CCaMK targets mediating LR formation. Constitutive and ectopic expression of *NIN* under the ubiquitin promoter resulted in a significant increase in LR number. We compared the responsiveness of two Rosaceae that have either retained *NIN* (*Dryas drummondii*) or lost it (*Fragaria vesca*) to stimulation with the constitutively active variant CCaMK^1−314^. Intriguingly, CCaMK^1−314^ was able to increase LR formation in *Dryas* but not in *Fragaria*, pointing to consequences of the evolutionary loss of *NIN* on root architecture. Taken together our data provide evidence for *NIN* as a molecular link between symbiosis-signaling and LR formation. Non-inoculated *nsp1* and *nsp2* mutant plants as well as HRLC of these mutants exhibited increased LR densities that were no further increased by expression of CCaMK^1-314^. We propose a model in which LR density is balanced by the activation of *NIN* expression by SymRK and CCaMK and the LR suppressing activity of NSP1 and NSP2.

## Introduction

### Arbuscular mycorrhiza fungi induces the formation of lateral roots

Root system architecture displays strong environment-responsive plasticity (Młodzińska-Michta, 2023). Most extant land plant species engage in symbiosis with arbuscular mycorrhiza (AM) fungi that deliver water and mineral nutrients to the host plant in return for lipids and carbohydrates (Parniske, 2008; Keymer *et al*., 2017; Luginbuehl & Oldroyd, 2017). Enhanced lateral root (LR) formation in response to inoculation with AM fungi was reported in monocots (e.g. rice [*Oryza sativa*] and maize [*Zea mays*] (Paszkowski & Boller, 2002; Gutjahr *et al*., 2009; Chiu *et al*., 2022)) and dicots (e.g. *Medicago truncatula* (Oláh *et al*., 2005; Chiu *et al*., 2022)). In this study, we aimed to elucidate the mechanistic connection between symbiosis-signaling components and the induction of LR development.

### LR and the root nodules of legumes are structurally related and share a common regulatory transcription factor, ASL18/LBD16

Lateral organs form on plant roots during the nitrogen-fixing root nodule symbiosis (RNS) and are called root nodules (RNs). Based on closely related structural features, such as the root and RN vasculature, RN organogenesis may have evolved by co-opting critical steps from the ancestral LR developmental program (Hirsch *et al*., 2001; Schiessl *et al*., 2019). This hypothesis predicts an overlap in regulatory and executive genes involved in the development of these two types of lateral organs. This concept received support from the discovery that RN development in the model legumes *Lotus japonicus* and *M. truncatula* requires a regulatory cascade controlled by the transcription factor Asymmetric Leaves 2-like 18 (ASL18, ortholog of Lateral organ Boundaries Domain 16 [LBD16] in *M. truncatula*) (Schiessl *et al*., 2019; Soyano *et al*., 2019), whose ortholog in Arabidopsis was described as being critical for LR development (Okushima *et al*., 2007; Goh *et al*., 2012, 2019).

### NIN is essential for nodule organogenesis and regulates *LBD16* expression

Exclusively in legumes, *LBD16a* harbors an intronic binding site for the transcription factor Nodule Inception (NIN) (Soyano *et al*., 2019) which regulates its expression to trigger nodule formation (Soyano *et al*., 2019; Shahan & Benfey, 2020). *NIN* is essential for RN organogenesis and rhizobial infection (Schauser *et al*., 1999) and encodes a transcriptional regulator (Soyano *et al*., 2015). *NIN* expression is itself modulated by a promoter which has in legumes increased its complexity and reaches up to 100 kb 5’ of the transcriptional start site (Liu *et al*., 2019a). *NIN* expression is regulated by a complex spatio-temporal network of regulatory inputs (Soyano *et al*., 2021), including the phytohormones cytokinin (Gonzalez-Rizzo *et al*., 2006; Heckmann *et al*., 2011; Liu *et al*., 2019a) and gibberellic acid (Takeda *et al*., 2015; Akamatsu *et al*., 2021) and symbiosis-induced signal transduction (Singh *et al*., 2014; Cathebras *et al*., 2022). Recently, the oscillatory nature of its expression pattern in synchronicity with NSP1 expression has been impressively documented (Soyano *et al*., 2024).

### Symbiosis-induced lateral root formation

The observation that both symbiotic lipo-chito-oligosaccharides (LCOs) (Oláh *et al*., 2005; Maillet *et al*., 2011; Chiu *et al*., 2022) and AM fungi (Paszkowski & Boller, 2002; Oláh *et al*., 2005; Gutjahr *et al*., 2009; Chiu *et al*., 2022) trigger an increase in LR density supports the idea that key regulatory signal transduction components shared between AM and RNS might be responsible for the increase of LR density. A set of so-called “common *Sym* genes” (Kistner & Parniske, 2002) – including, among other genes, *SymRK*, *CCaMK* and *Cyclops* – is defined by their mutant phenotype; a loss of both AM and RNS (Stracke *et al*., 2002; Lévy *et al*., 2004; Mitra *et al*., 2004; Tirichine *et al*., 2006; Yano *et al*., 2008).

Symbiosis-induced signal transduction requires *Symbiosis Receptor-like Kinase* (*SymRK* (Stracke *et al*., 2002)) and affects *NIN* transcription *via* a complex formed by the Calcium Calmodulin-dependent Protein Kinase (CCaMK (Lévy *et al*., 2004; Mitra *et al*., 2004; Tirichine *et al*., 2006)) and its phosphorylation target Cyclops (Messinese *et al*., 2007; Yano *et al*., 2008; Singh *et al*., 2014).

CCaMK/Cyclops may assemble into larger complexes with, among others, the GRAS proteins DELLA (Pimprikar *et al*., 2016), Nodulation Signalling Pathway 1 (NSP1) and NSP2 (Catoira *et al*., 2000; Oldroyd & Long, 2003; Kaló *et al*., 2005; Smit *et al*., 2005; Heckmann *et al*., 2006; Murakami *et al*., 2006), and/or the Interacting Protein of NSP2 (IPN2) (Kang *et al*., 2014; Xiao *et al*., 2020).

### GRAS proteins are involved in root endosymbiosis and the regulation of strigolactone biosynthesis

*NSP1* and *NSP2* are both indispensable for RNS and are involved in LCO-mediated signaling in RNS and AM (Catoira *et al*., 2000; Oldroyd & Long, 2003; Smit *et al*., 2005; Hirsch *et al*., 2009; Maillet *et al*., 2011; Delaux *et al*., 2013; Nagae *et al*., 2014; Jin *et al*., 2016). Importantly, and potentially epistatic to all other phenotypes, NSPs regulate the expression of genes involved in the biosynthesis of strigolactones during phosphate starvation in *M. truncatula* (Li *et al*., 2022a), barley (*Hordeum vulgare*) (Li *et al*., 2022a) and rice (Liu *et al*., 2011; Yuan *et al*., 2023). NSP1 and NSP2 form a complex that has been described to interact with DELLA (Jin *et al*., 2016) and IPN2, a MYB transcription factor that was reported to regulate *NIN* expression presumably *via* direct binding to its promoter (Xiao *et al*., 2020).

### Ectopic *SymRK* expression and deregulated CCaMK and Cyclops versions lead to spontaneous root nodule development

Ectopic overexpression of *SymRK* (Ried *et al*., 2014) or its intracellular domain (Saha *et al*., 2014), or auto-active versions of CCaMK (CCaMK^1-314^ and CCaMK^T265D^)^26,44,45^ (Gleason *et al*., 2006; Tirichine *et al*., 2006; Takeda *et al*., 2012) or auto-active Cyclops (Cyclops^S50D, S154D^ hereafter called Cyclops^DD^)(Singh *et al*., 2014) can spontaneously activate (i.e. in the absence of rhizobia or AM fungi) symbiosis-related gene expression as well as RN organogenesis in a *NIN* and *NSP1*/*NSP2* dependent manner (Hayashi *et al*., 2010; Madsen *et al*., 2010; Singh *et al*., 2014).

### Aim of this study

The identification of LBD16a/ASL18a provided a key connection between LR and RN initiation. A role of *NIN* and its upstream regulators for rhizobial infection and for RN development are supported by a large body of evidence. However, a role of common symbiosis gene and *NIN* in LR initiation has remained speculative, as genetic evidence for their capacity to induce LR development has been lacking.

### Advantages of a lateral root assay in liquid medium and in the absence of a shoot

Regulatory networks dictating the “LR-to-RN” decision are complex and remain only partially understood. Previous studies were plagued by the multidimensional impact of the shoot on the LR versus RN decision, whose pronounced regulatory impact masks the quantitative contributions of root-endogenous regulatory networks. In order to abolish the influence of the shoot thus permitting an assessment of the direct impact of common symbiosis genes on LR induction, we established a hairy root liquid culture (HRLC) system. Root organ cultures are unable to support rhizobia induced RN development (Raggio *et al*., 1957; Tsikou *et al*., 2018), therefore providing an elegant tool to uncouple lateral organ development from systemic control via the rhizobia- and nitrate-dependent miR2111/TML regulon (Magori *et al*., 2009; Takahara *et al*., 2013; Tsikou *et al*., 2018; Sexauer *et al*., 2023) or the LR clock, which is driven by photosynthetic sucrose (Xuan *et al*., 2020; Kircher & Schopfer, 2023). Additionally, this system allowed for the determination of the quantitative impact of transgenes on LR formation without the impact of nutrient gradients or substrate particles. Unlike solid media, which can introduce nutrient heterogeneity due to nutrient adsorption to substrate particles and thereby create localized LR-inducing hotspots – or even trigger LR formation through physical contact – liquid conditions provide a chemically and mechanically more homogeneous environment. By removing shoot-derived input, nutrient gradients and substrate-mediated physical stimuli, this HRLC system enabled a directly comparable quantitative assessment of LR induction by genes involved in symbiosis-induced signal transduction.

## Results

### Establishment of a hairy root culture in liquid medium

We found that ectopic and constitutive expression of *Cyclops^DD^*under the control of the *L. japonicus Ubiquitin* promoter in transgenic roots of composite *L. japonicus* plants leads to a drastic increase of LR numbers (Fig. **S1**). This observation motivated us to examine whether spontaneous activation of symbiosis signaling – mediated by auto-active versions of common symbiosis genes such as *Cyclops^DD^* – induce an increase of LRs in *L. japonicus*. However, the induction of spontaneous RNs on the same root system and the consequential inhomogeneity of the readout in terms of type of LR organ motivated us to develop a quantitative LR assay with shoot-less hairy root systems grown in liquid cultures (Fig. **S2**). All transgenic roots expressed free GFP as a visual transformation marker. To standardize the assay, root sections of equal length (1.5 cm, always including the root tip) were excised from fluorescent roots at four weeks post inoculation with *Agrobacterium rhizogenes* and grown individually in liquid medium (Fig. **S2**).

### Deregulated variants of common symbiosis genes stimulate lateral root formation in root liquid cultures

We generated transgenic *L. japonicus* hairy root liquid cultures (HRLC) transformed with an empty vector control (EV) or with a construct ectopically expressing either *SymRK*, *CCaMK^1−314^*, *CCaMK^T265D^*, or *Cyclops^DD^* under the control of the *L. japonicus Ubiquitin* promoter (*Ubq_pro_* ^56^). Notably, all four constructs significantly increased LR number relative to the EV (Fig. **1a, b**). Specifically, LR number increased by approximately 75% with the ectopic expression of *SymRK* and by 110% with *CCaMK^1−314^*, *CCaMK^T265D^*, or *Cyclops^DD^*at 10 days of incubation (doi) in HRLC, in comparison to control HRLC transformed with the EV (Fig. **1a**). HRLC ectopically expressing Cyclops in which the serine residues S50 and S154 were replaced with phospho-ablative alanine (A) (*Cyclops^AA^*), however, displayed a similar number of LRs as HRLC transformed with the EV (Fig. **S3**). Additionally, the significantly higher numbers of LRs mediated by the ectopic overexpression of *SymRK*, *CCaMK^1−314^*, *CCaMK^T265D^* or *Cyclops^DD^* were maintained over a time course of 30 days (Fig. **S4**). The observation that in *L. japonicus*, LR number can be increased by ectopically overexpressing *SymRK, Cyclops^DD^*, *CCaMK^1-314^*or *CCaMK^T265D^*, reveals a function of these common symbiosis genes in the regulation of LR formation.

**Figure 1.**
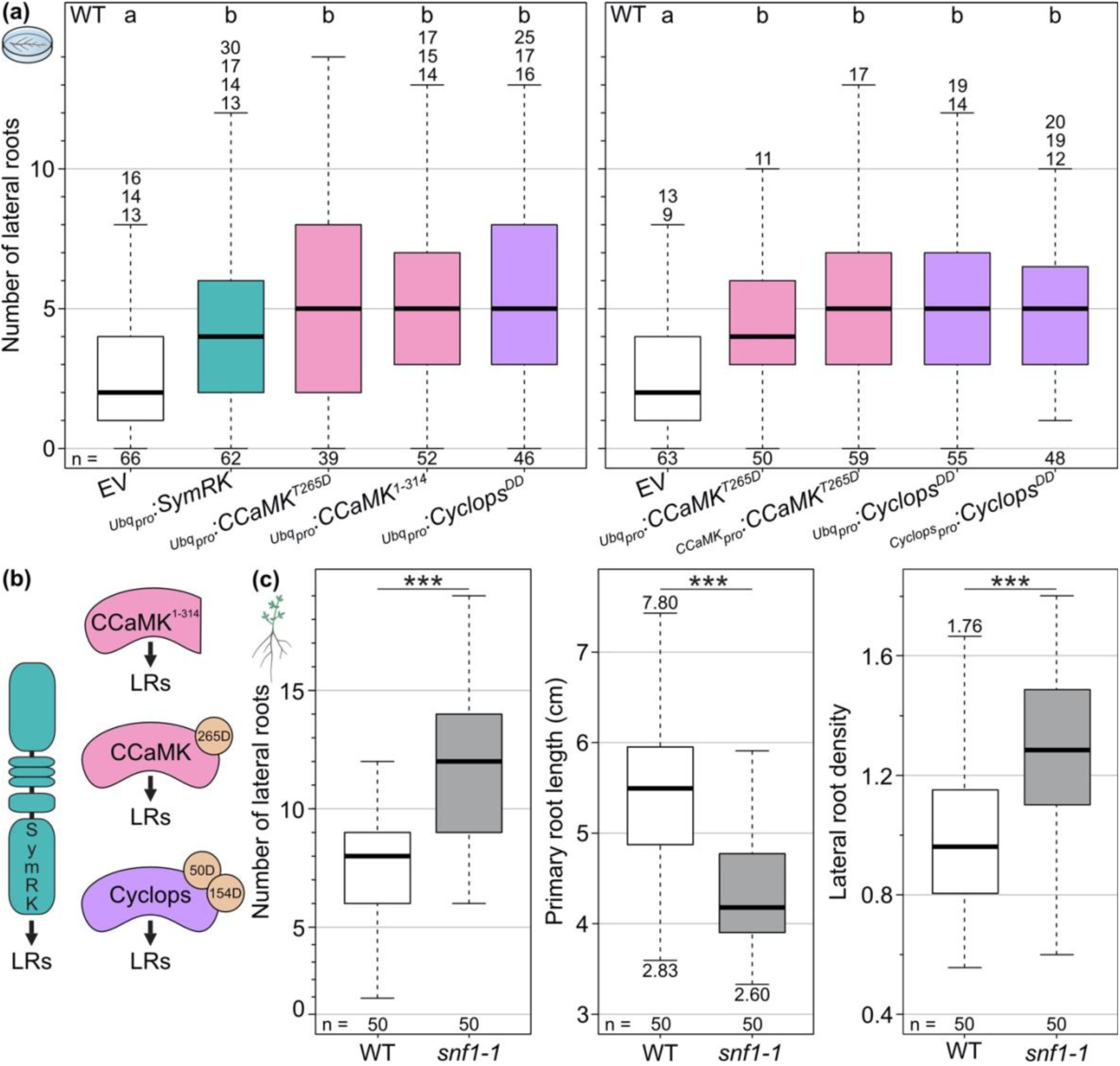
Deregulated versions of CCaMK or Cyclops or ectopic overexpression of *SymRK* stimulate lateral root formation. **(a)** Hairy root liquid cultures of *L. japonicus* WT transformed with the empty vector (EV), *Ubq_pro_:SymRK*, *Ubq_pro_:CCaMK^T265D^*, *Ubq_pro_:CCaMK^1-314^* or *Ubq_pro_:Cyclops^DD^*(**left panel**), or with the empty vector (EV), *Ubq_pro_:CCaMK^T265D^*, *CCaMK_pro_:CCaMK^T265D^*, *Ubq_pro_:Cyclops^DD^*or *Cyclops_pro_:Cyclops^DD^* (**right panel**) were incubated in MSR liquid medium. Box plots represent the number of lateral roots per root culture after 10 days of incubation. Data were subjected to a Kruskal-Wallis test followed by a Dunn’s *post hoc* analysis; *p* < 0.05. **(b) Model:** Auto-active versions of CCaMK or Cyclops or ectopic overexpression of *SymRK* increase lateral root numbers **(c) *L. japonicus* WT and *snf1-1* plants** (*snf1-1* carries a point mutation leading to the amino acid replacement CCaMK^T265I^) grown for 30 days. Box plots represent the number of lateral roots, lateral root density or primary root length. Note that while the primary root length of *snf1-1* plants is decreased, lateral root number and density are significantly increased. Data were subjected to Welch’s *t*-test; \*\*\**p* < 0.001. Lateral root density: number of lateral roots per cm of the primary root. n: number of roots/plants analyzed.

To analyze to what extent the chosen promoter was relevant for the increased LR numbers, we transformed *L. japonicus* WT roots with *Cyclops^DD^* or *CCaMK^T265D^* driven either by their own promoter (*Cyclops_pro_:Cyclops^DD^* or *CCaMK_pro_:CCaMK^T265D^*) or driven by the *Ubiquitin* promoter. We observed that these two auto-active versions stimulated LR formation to the same extent regardless of the promoter (Fig. **1a, b**).

### The *snf1-1* mutant encoding CCaMK^T265I^ produces more lateral roots

In order to test to what extend the changes imposed by the HRLC contribute to these effects, we tested the *L. japonicus* mutant line *spontaneous nodule formation 1-1* (*snf1-1*) which carries the CCaMK^T265I^ mutant version (Tirichine *et al*., 2006) at the same position as the previously tested T265D replacement. The important difference was that the availability of this line allowed us to test the impact of this auto-active version of CCaMK in an intact plant assay without the involvement of transgenes or HRLC. On intact non-inoculated plants, we observed that the *snf1-1* line displayed a higher LR density and more LRs compared to WT plants (Fig. **1c**). This increase in LRs by a) CCaMK^T265I^ in a whole plant setting (Fig. **1c**) and b) CCaMK^T265D^ in HRLC expressed under its own promoter (Fig. **1a**) provide strong and independent evidence that auto-active versions of CCaMK increase the number of LRs.

### Hairy roots without shoot exclusively produced lateral roots and no nodules or nodule-like structures

Importantly, none of the transgenes, previously shown to spontaneously activate RN organogenesis when expressed in hairy roots of intact *L. japonicus* plants (Gleason *et al*., 2006; Tirichine *et al*., 2006; Takeda *et al*., 2012; Ried *et al*., 2014; Singh *et al*., 2014), induced spontaneous RNs on any of the 2063 HRLCs generated throughout the course of this study. The complete absence of spontaneous RNs suggests that shoot-derived signals are required to permit not only rhizobia-induced RN development as described earlier (Raggio *et al*., 1957; Tsikou *et al*., 2018; Li *et al*., 2022b), but also spontaneous RN development in *L. japonicus.* Taken together, our results indicate that the expression of the deregulated versions CCaMK^T265D^, CCaMK^1-314^ or Cyclops^DD^, or the ectopic over-expression of *SymRK*, results in the stimulation of the LR developmental program (Fig. **1**).

### *NIN* is required for the increase in lateral root number stimulated by deregulated CCaMK

*NIN* is indispensable for spontaneous RN formation mediated by deregulated variants of CCaMK (Marsh *et al*., 2007; Madsen *et al*., 2010) or Cyclops (Singh *et al*., 2014). To examine whether *NIN* was required for auto-active CCaMK-mediated increase of LR numbers, we transformed WT and *nin-2* mutant roots with *Ubq_pro_:CCaMK^1−314^*. Intriguingly, ectopic expression of *CCaMK^1−314^* in *nin-2* HRLC led to a reduction in LR number, in comparison to HRLC transformed with the EV or with *Ubq_pro_:CCaMK* (Fig. **2a**). HRLC of WT transformed with *Ubq_pro_:CCaMK^1−314^* formed, in the same experiment, significantly more LRs (Fig. **2a**), consistent with previous results (Fig. **1a**). It is tempting to speculate that CCaMK^1−314^ activates a repressor of LR formation that is not only counteracted but outcompeted by NIN in the WT (Fig. **2a, d**). Importantly, this reduction in LR number in HRLC of *nin-2* was dependent on the ectopic expression of *CCaMK^1−314^*, as LR density and primary root length was comparable in the non-nodulating *nin-2* mutant and WT plants (Fig. **2c**).

**Figure 2.**
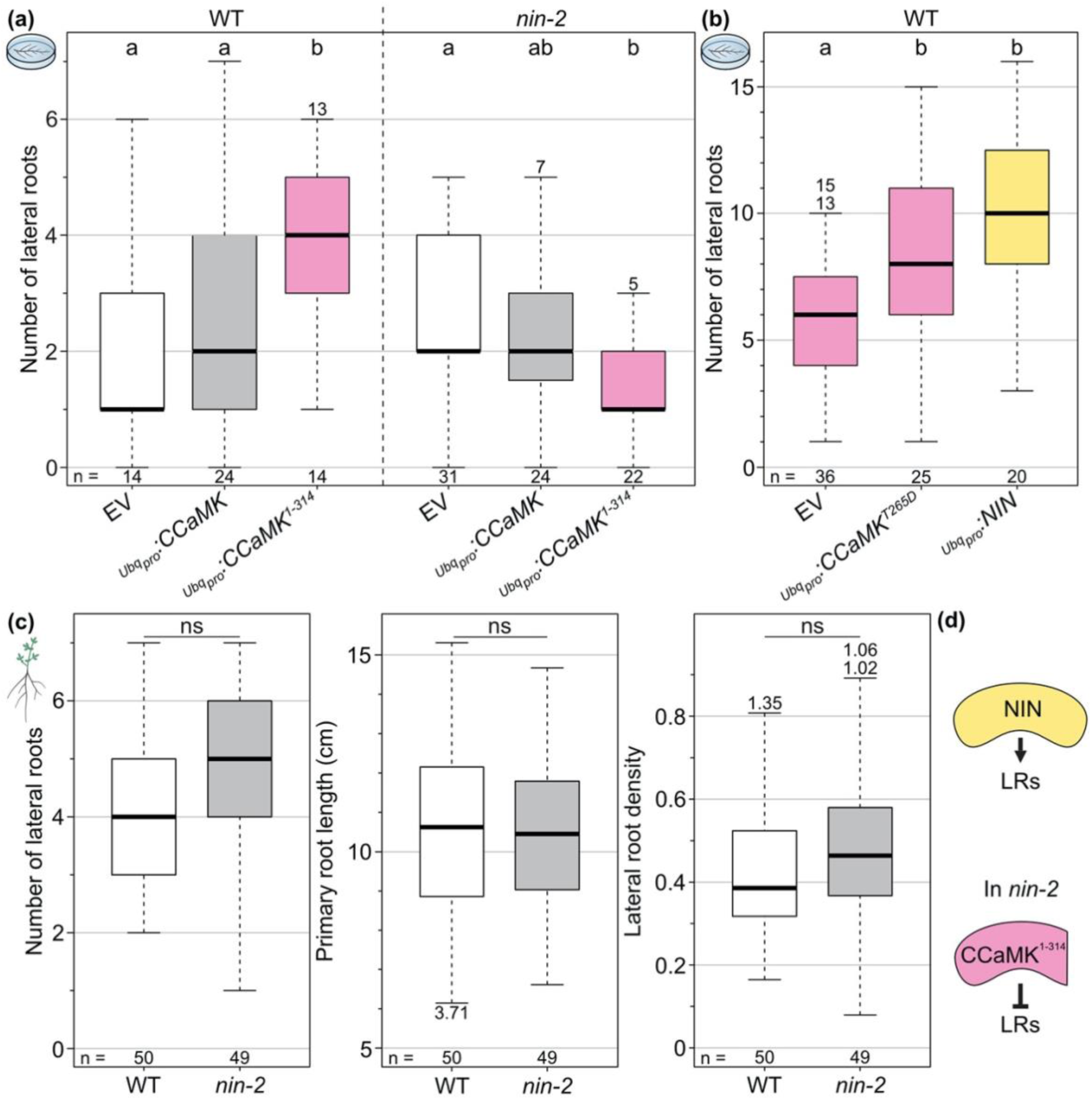
Ectopic overexpression of *NIN* increases lateral root numbers, while ectopic overexpression of *CCaMK^1-314^* in the *nin-2* background decreases lateral root numbers. **(a-b)** Hairy root liquid cultures of (a) *L. japonicus* WT and *nin-2* transformed with the empty vector (EV), *Ubq_pro_:CCaMK* or *Ubq_pro_:CCaMK^1-314^* and (**b**) *L. japonicus* WT transformed with the empty vector (EV), *Ubq_pro_:CCaMK^T265D^* or *Ubq_pro_:NIN*. Box plots represent the number of lateral roots per root culture after 10 days of incubation. Data were subjected to a Kruskal-Wallis test followed by Dunn’s *post hoc* analysis; *p* < 0.05. (c) *L. japonicus* WT and *nin-2* plants grown for 30 days. Box plots represent the number of lateral roots, primary root length and lateral root density. Data were subjected to Welch’s *t*-test; \**p* < 0.05; \*\**p* < 0.01; \*\*\**p* < 0.001. ns: not significant. **(d) Model:** Ectopic overexpression of *NIN* increases lateral root numbers, while ectopic overexpression of *CCaMK^1-314^*in a *nin-2* background decreases lateral root numbers. Lateral root density: number of lateral roots per cm of the primary root. n: number of roots/plants analyzed.

### Ectopic expression of *NIN* stimulates lateral root formation

Ectopic expression of *NIN* was found to induce the formation of enlarged bumps and malformed LRs in the absence of a symbiont in composite plants of *L. japonicus* (Soyano *et al*., 2013) and soybean (*Glycine max*)(Fu *et al*., 2022). However, increased LR numbers were not reported. We thus asked whether ectopic expression of *NIN* stimulates the formation of LRs in HRLC, which are unable to support spontaneous RN development. At 10 doi, we observed a significant increase in LR number in WT HRLC transformed with *Ubq_pro_:CCaMK^T265D^*or *Ubq_pro_:NIN* in comparison to EV controls (Fig. **2b**). These results indicate that ectopic expression of the transcriptional regulator gene *NIN* is sufficient to trigger the development of more LRs (Fig. **2**). In a subset (∼12%) of the HRLC ectopically expressing *NIN*, roots curled and formed spirals (Fig. **S5**), an unexplained phenomenon somewhat reminiscent of the spiral roots observed by Schiessl *et al*. (Schiessl *et al*., 2019) upon ectopic expression of *LBD16* in *M. truncatula*. We interpret the low percentage of ∼12% such that in these rare cases, the expression pattern of *NIN* is dictated by the genome environment of the T-DNA insertion. This phenomenon was never observed with any of the other transgenes tested in this work that led to an increase in LR numbers in HRLC (**Figs. 1-5**) and are considered inducers of *NIN* expression. This exclusive occurrence suggests a restrictive expression pattern dictated by the endogenous *NIN* promoter that does not allow root spiraling.

### Auto-active CCaMK^T265D^ stimulates lateral root formation in the absence of *Cyclops*

Cyclops is essential for infection thread formation but is dispensable for rhizobia or CCaMK^T265D^-induced nodule organogenesis (Yano *et al*., 2008). To elucidate whether *Cyclops* is dispensable for the stimulation of LR formation mediated by deregulated CCaMK, we ectopically expressed *CCaMK^T265D^* in roots of the *cyclops-3* mutant in HRLC (Fig. **3a**). Similar to WT HRLC, *cyclops-3* HRLC ectopically expressing *CCaMK^T265D^* produced significantly more LRs in comparison to the HRLC transformed with the EV (Fig. **3**).

This finding, together with the observation that Cyclops^DD^ in the *ccamk-3* background leads to increased LR formation, highlights the existence of signaling components downstream of *CCaMK* at the same hierarchical level as *Cyclops* that can stimulate LR development in the absence of *Cyclops* (Fig. **3**).

**Figure 3.**
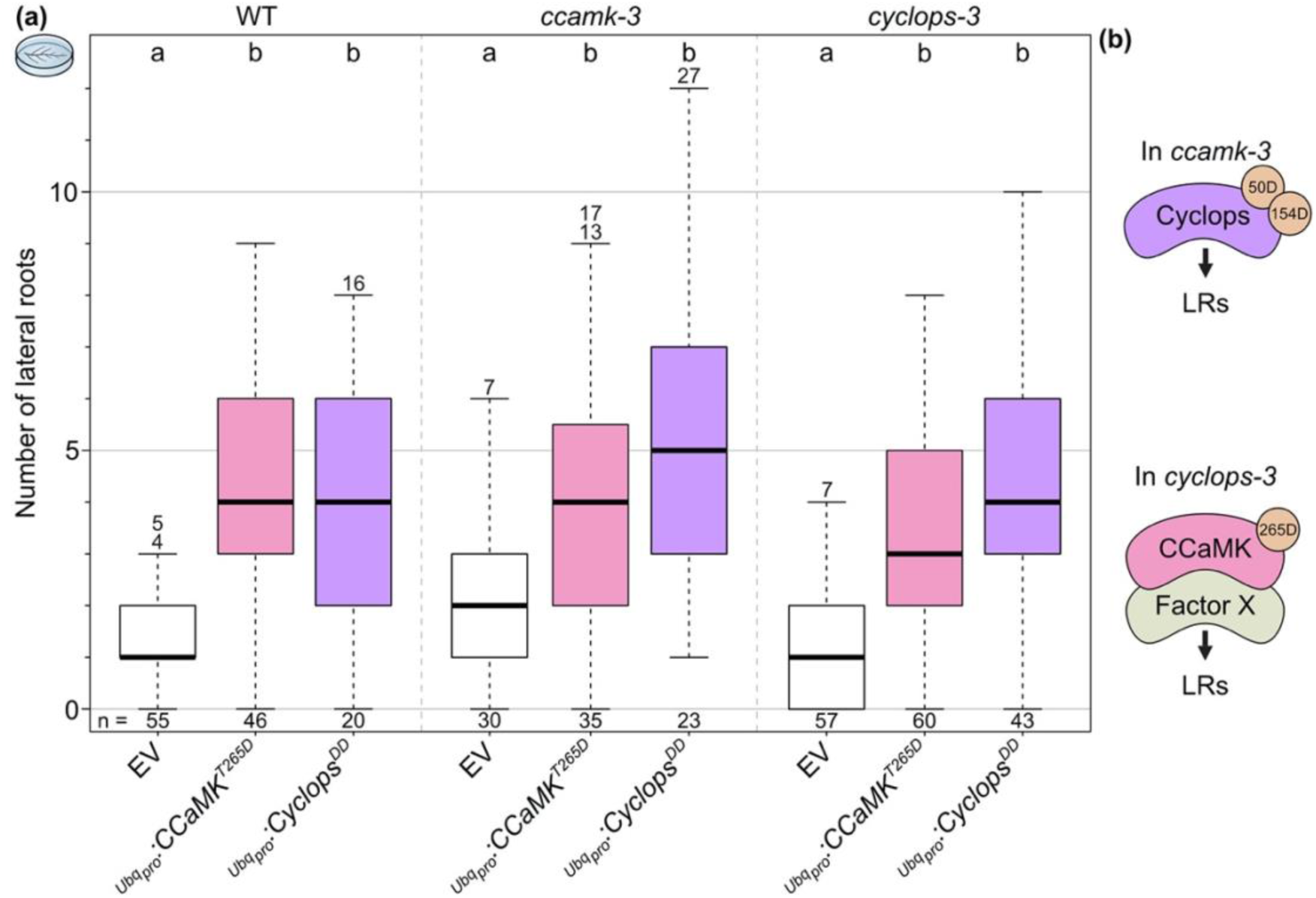
Increased lateral root numbers mediated by deregulated versions of CCaMK or Cyclops are not dependent on endogenous *CCaMK* or *Cyclops*. **(a)** Hairy root liquid cultures of *L. japonicus* WT, *ccamk-3* and *cyclops-3* transformed with the empty vector (EV), *Ubq_pro_:CCaMK^T265D^* or *Ubq_pro_:Cyclops^DD^*. Box plots represent the number of lateral roots per root culture after 10 days of incubation. Data were subjected to a Kruskal-Wallis test followed by Dunn’s *post hoc* analysis; *p* < 0.05. **(b) Model:** Expression of Cyclops^DD^ increases lateral root numbers in the *ccamk-3* background. Expression of CCaMK^T265D^ increases lateral root numbers also in the absence of *Cyclops* indicating the existence of a CCaMK^T265D^ target that acts redundantly with Cyclops (Factor X). n: number of roots analyzed.

### *NSP1* and *NSP2* are potential repressors of lateral root formation mediated by NIN

Potential candidates for signaling components downstream of CCaMK may be NSP1 and NSP2, as they have been placed into a conceptual regulatory complex involving CCaMK, Cyclops, DELLA and IPN2 controlling *NIN* expression (Jin *et al*., 2016). HRLC of the *nsp1-1* or *nsp2-2* mutants formed significantly more LRs than those of the WT transformed with the EV control, with no further increase when either mutant was transformed with *Ubq_pro_:CCaMK^1^*^−*314*^ (Fig. **4b, c**). Additionally, both *nsp* mutant plants displayed a higher LR density than WT plants (Fig. **4a**). Taken together, NSP1 and NSP2 appear to be negative regulators of LR development. Whether this effect is due to their role in regulating *NIN* expression or their positive effect on the expression of strigolactone biosynthesis genes or a combination of both, requires further investigation.

**Figure 4.**
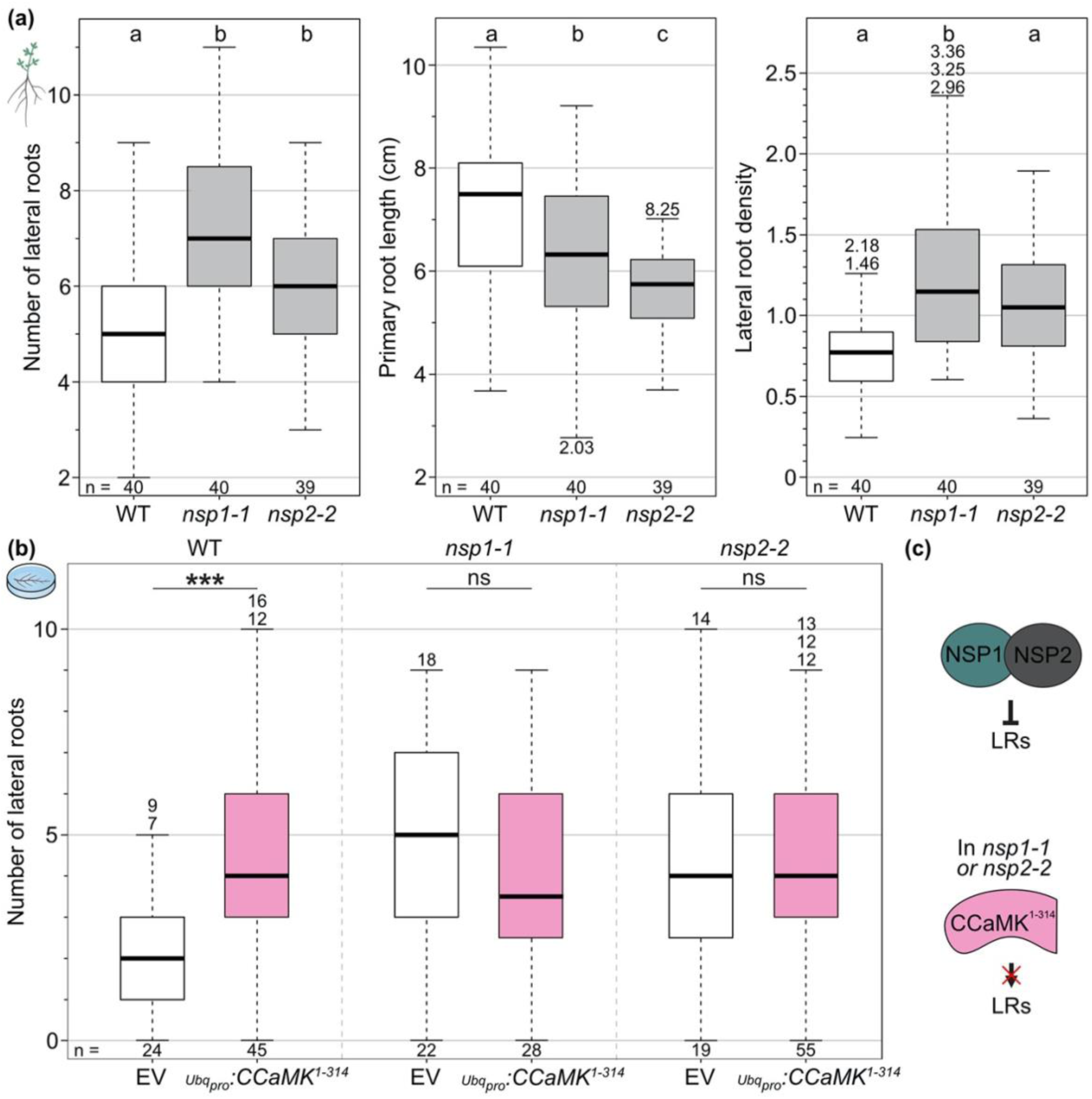
*Lotus japonicus nsp1-1* and *nsp2-1* mutant plants feature shorter primary roots and increased lateral root numbers. **(a) *L. japonicus* WT, *nsp1-1* and *nsp2-2* plants** grown for 30 days. Box plots represent the number of lateral roots, lateral root density and primary root length. Data were subjected to a Kruskal-Wallis test followed by Dunn’s *post hoc* analysis; *p* < 0.05. **(b) Hairy root liquid cultures** of *L. japonicus* WT, *nsp1-1* and *nsp2-2* transformed with the empty vector (EV) or *Ubq_pro_:CCaMK^1-314^*. Box plots represent the number of lateral roots per root culture after 14 days of incubation. Data were subjected to Welch’s *t*-test; \*\*\**p* < 0.001. ns: not significant. (c) Model: *L. japonicus nsp1-1* and *nsp2-1* mutants formed more lateral roots than the WT, suggesting that NSP1 and NSP2 negatively regulate lateral root development. Ectopic expression of *CCaMK^1-314^* in the *nsp1-1* or *nsp2-1* background did not increase lateral root numbers further. Lateral root density: number of lateral roots per cm of the primary root. n: number of roots/plants analyzed.

### Lateral root induction mediated by deregulated CCaMK is conserved in *Dryas drummondii* but not in *Fragaria vesca*

Our data revealed *NIN* as a main player in the formation of LRs. As *NIN* has been lost multiple times within the Fabales, Fagales, Cucurbitales, and Rosales (FaFaCuRo) clade (Soltis *et al*., 1995; Doyle, 2011; Griesmann *et al*., 2018), the question arises whether the loss of *NIN* during evolution had consequences on root development. We tested the LR inducing capability of *LjCCaMK^1^*^−*314*^ in two members of the Rosaceae, the actinorhizal plant *Dryas drummondii*, which carries a *NIN* ortholog, and in *Fragaria vesca,* a non-nodulating species that lost a functional *NIN* (Griesmann *et al*., 2018). We generated composite plants with transgenic roots and observed a significant increase in LR number and consequently in LR density on *D. drummondii* roots transformed with *Ubq_pro_:LjCCaMK^1−314^* in comparison to roots transformed with the EV or with *Ubq_pro_:Myc-CCaMK* (Fig. **5a**). By contrast, *F. vesca* roots transformed with *Ubq_pro_:LjCCaMK^1−314^*did not display any significant increase in LR number or primary root length and thus in LR density in comparison to roots transformed with the control constructs (Fig. **5b**). Taken together, these results indicate that the induction of LRs mediated by CCaMK^1−314^ is not specific to legumes (Fabales) and suggest that *NIN* is required to mediate this developmental response in species outside of the Fabales.

**Figure 5.**
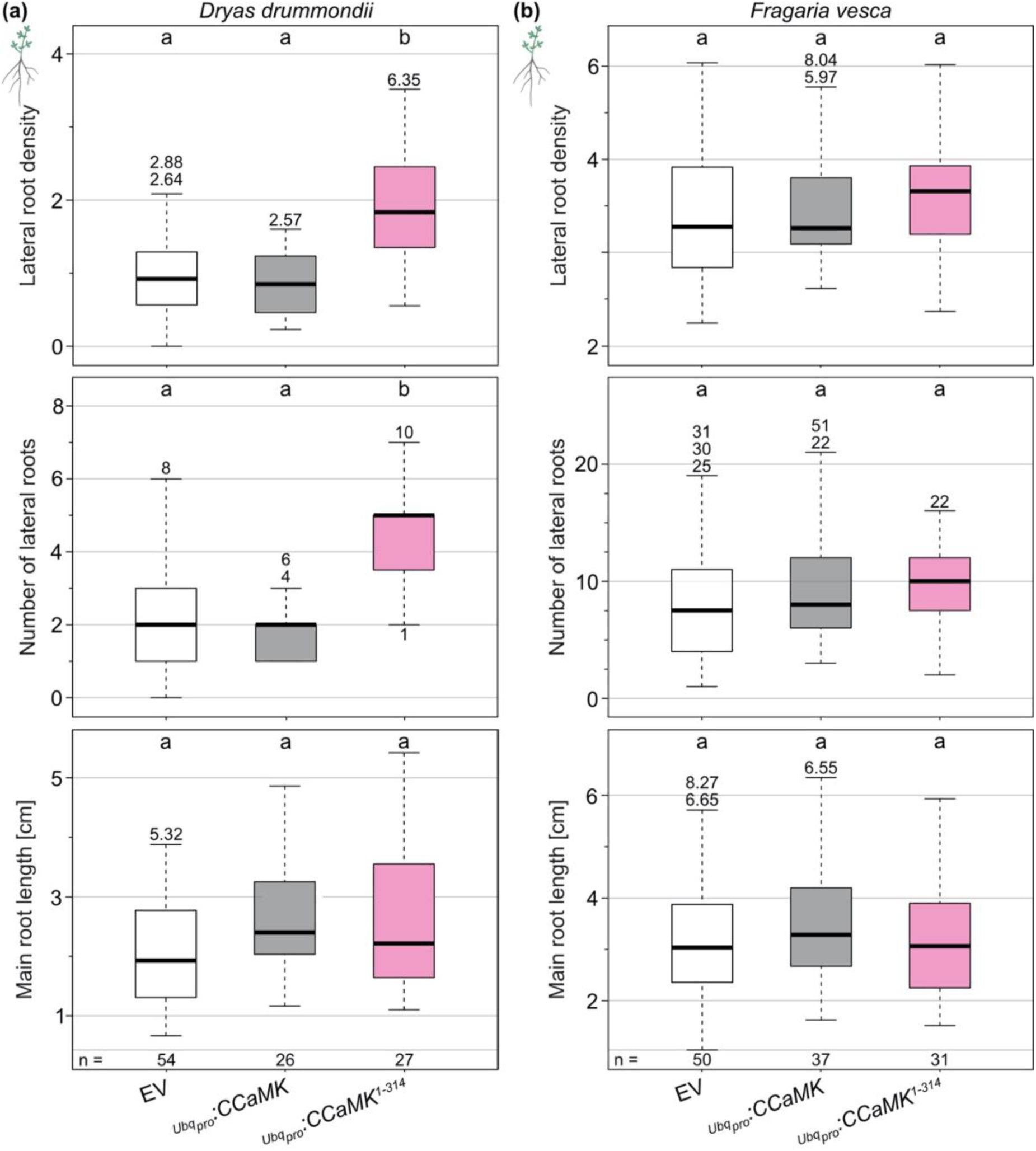
Ectopic expression of *LjCCaMK^1-314^* increases lateral root density in *Dryas drummondii* but not significantly in *Fragaria vesca*. **(a)** *D. drummondii* and **(b)** *F. vesca* composite plants each bearing a single hairy root (hereafter called “main root”) transformed with the empty vector (EV), *Ubq_pro_:LjCCaMK* or *Ubq_pro_:LjCCaMK^1-314^* were transferred to Weck jars 45 and 42 days post *Agrobacterium rhizogenes* inoculation, respectively. Box plots represent the lateral root density (upper panel), the number of lateral roots (middle panel) and the main root length (lower panel) at 65 days post *Agrobacterium rhizogenes* inoculation. Data were subjected to Kruskal-Wallis test followed by Dunn’s *post hoc* analysis; *p* < 0.05. Lateral root density: number of lateral roots per cm of the main root. n: number of plants analyzed.

## Discussion

### Ectopic expression of the common symbiosis gene *SymRK*, and auto-active versions of CCaMK and Cyclops increase LR number

In this study, we report a novel regulatory role of the symbiosis related SymRK, CCaMK and Cyclops and NIN for the development of LRs. We hypothesize that the LR induction by the deregulated versions of CCaMK and Cyclops likely works *via* the transcriptional activation of *NIN*. The existence of this connection between the regulation of *NIN* and LR formation was significantly but not exclusively supported by the use of HRLC in which the initiation of lateral organs from the root are limited to LRs.

We found that ectopic overexpression of *SymRK* or expression of *CCaMK^T265D^* or *Cyclops^DD^* driven by their native promoters in the absence of external symbiotic stimuli significantly increased LR formation (Fig. **1a, b**). Furthermore, expression of *CCaMK^T265D^* in the *cyclops-3* mutant also resulted in an increase in LR number (Fig. **3**), suggesting the existence of genetic redundancy at the hierarchical level of Cyclops in the context of symbiosis-associated LR development (Fig. **6**). A strong candidate for a partly redundant factor is ERN1 which together with Cyclops coordinates *NIN* expression, and a *cyclops-3 ern1-1* double mutant did not produce any white or pink nodules upon *Mesorhizobium loti* inoculation whereas single mutants produced white nodules (Liu *et al*., 2019b). In addition, ERN1 is required for the formation of spontaneous nodule mediated by auto-active CCaMK^T265D^ (Kawaharada *et al*., 2017).

Importantly, we never observed to formation of spontaneous RNs in any of the HRLC initiated in this work. Although HRLC have been previously reported to be unable to support rhizobium induced RN development (Tsikou *et al*., 2018), here we reveal that the ability to form the rhizobium signal independent (aka spontaneous) RN organ *per se*, is dependent on signals from the shoot. HRLC therefore provides an ideal platform to identify missing shoot-derived factors required for RN initiation and or development.

Compared to WT plants, LR density was significantly higher in *snf1-1* mutant plants, carrying a point mutation leading to the amino acid replacement CCaMK^T265I^, confirming the role of CCaMK in LR regulation also in the whole plant context. This important observation rules out the possibility that, at least for CCaMK, the observed increase in LRs was a consequence of the absence of the shoot (Fig. **1c**).

**Figure 6.**
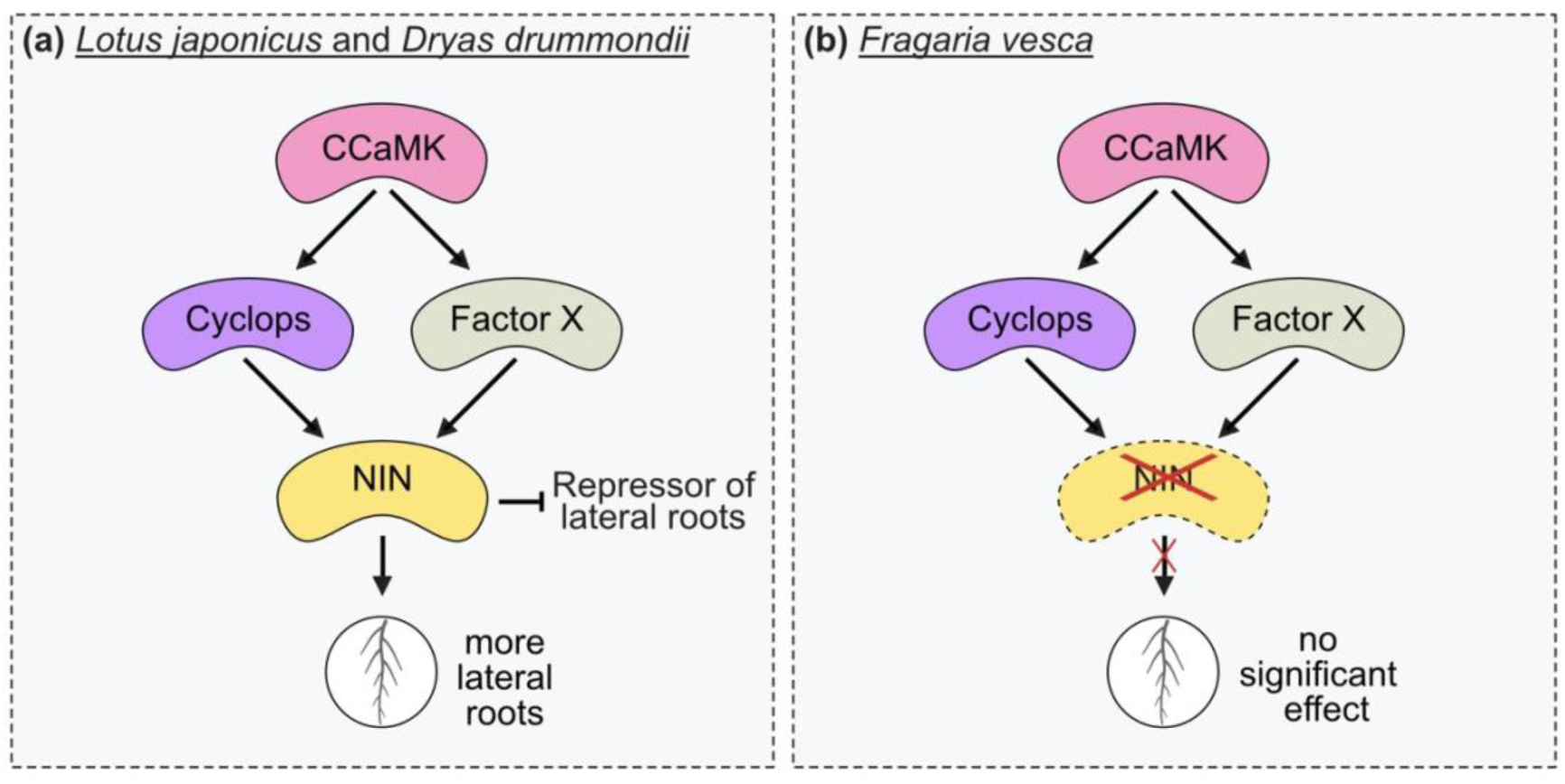
A model for the regulation of lateral root development promoted *via* the transcriptional activation of *NIN* and antagonized by a – CCaMK activated – “Repressor of lateral roots”. **(a)** In *L. japonicus* and *D. drummondii,* CCaMK induce the formation of lateral roots *via* the activation of Cyclops and/or Factor X that in turn *trans*-activate *NIN* expression. *NIN* stimulates the formation of lateral roots. NIN induced by CCaMK might regulate/counteract the inhibition of lateral root formation mediated by a – CCaMK activated – “Repressor of lateral roots”. **(b)** In *F. vesca* that lost *NIN* from its genome, no CCaMK^1-314^-mediated induction of lateral root formation is observed.

Taken together with their previously described sufficiency in inducing spontaneous RNs (Gleason *et al*., 2006; Tirichine *et al*., 2006; Ried *et al*., 2014; Saha *et al*., 2014; Singh *et al*., 2014), our data demonstrate that the tested deregulated common symbiosis gene variants are sufficient for inducing LR development (Fig. **6**).

### *NIN* orchestrates lateral root development

*NIN*, a target of the CCaMK/Cyclops complex, is an essential regulator of nodule organogenesis (Schauser *et al*., 1999; Marsh *et al*., 2007; Singh *et al*., 2014). We found that ectopic overexpression of *NIN* in hairy root liquid cultures is sufficient to induce the formation of LRs that look anatomically similar to those formed in roots transformed with an EV (Figs. **2b**, **S5b**). Ectopic overexpression of *NIN* in transgenic roots of composite plants ectopically (over)expressing *NIN* was reported to lead to the spontaneous (in the absence of rhizobia) formation of LR organs that were either categorized as nodule primordium-like structures and LRs with enlarged tips in *L. japonicus* (Soyano *et al*., 2013) or spontaneous RN in soybean (Fu *et al*., 2022) and *M. truncatula* (Vernié *et al*., 2015). However, **an increase in LR number by ectopically overexpressing *NIN* has not been reported in legumes**. Considering that the absence of a shoot in our hairy root culture system is the main difference relative to previously used composite plant systems, this support the idea that the shoot-to-root communication is the key in this decision.

Interestingly, ectopic overexpression of a *NIN* paralog, which in contrast to legume NIN retained nitrogen responsiveness, in hairy roots of composite poplar (*Populus* sp., Maphigiales) plants led to an increase in LR density (Irving *et al*., 2022). This difference between the tested legumes and poplar may reflect evolutionary innovations specific to legumes in rhizobia- and nitrate-mediated root-to-shoot communication to regulate RNS (Magori *et al*., 2009; Takahara *et al*., 2013; Tsikou *et al*., 2018; Sexauer *et al*., 2023). Deciphering the regulatory mechanisms that decide whether a RN or a LR is formed upon appropriate *NIN* expression will require future research.

### NIN-regulated genes involved in local regulation of lateral organ formation

Our data raise the question as to how exactly *NIN* influences LR formation. There are multiple genes described to be directly or indirectly regulated by NIN that have the potential to influence LR development. These genes include the cell division–associated nuclear factor genes *NF-YA1* and *NF-YB1* whom expression is regulated by NIN via direct binding to their promoters (Soyano *et al*., 2013). In *L. japonicus* overexpression of *NF-YA1* or co-overexpression of *NF-YA1* and *NF-YB1* was sufficient to increase LR density (Soyano *et al*., 2019) and in Arabidopsis, expression of a variant of *NF-YA2* resistant to the regulation by microRNA169 increased LR density (Sorin *et al*., 2014). It is therefore tempting to speculate that ectopic expression of *NIN* activates *NF-Y* expression thereby inducing LR formation, likely also involving *ASL18a* (Soyano *et al*., 2019). NIN also regulates the expression of the cytokinin receptor Cytokinin Response 1 (CRE1) (Vernié *et al*., 2015) which negatively interferes with LR development (Gonzalez-Rizzo *et al*., 2006; Laffont *et al*., 2015; Herrbach *et al*., 2017).

### NIN-regulated genes involved in systemic regulation of lateral organ formation

Additional candidates are the *NIN*-regulated genes encoding the root-to-shoot signaling peptides Clavata/Embryo Surrounding Region-Related Root Signal 1 (CLE-RS1) and CLE-RS2 (Soyano *et al*., 2014). The cognate receptors of CLE peptides and CEPs, *Lj*HAR1 (Wopereis *et al*., 2000; Krusell *et al*., 2002; Nishimura *et al*., 2002; Okamoto *et al*., 2013) and *Mt*CRA2 (Huault *et al*., 2014; Gautrat *et al*., 2020), respectively, are also implicated in the regulation of root system architecture. While *Lj*HAR1 regulates root morphology locally as well as systemically (Hayashi-Tsugane & Kawaguchi, 2022), *Mt*CRA2 controls LR formation locally *via Mt*CEP1 (Imin *et al*., 2013; Huault *et al*., 2014; Mohd-Radzman *et al*., 2016).

### The interplay between NIN and NLPs might play a role in the regulation of lateral root formation

NIN and NIN-like proteins (NLPs) physically interact and can form homo- and hetero-complexes (Lin *et al*., 2018) that bind competitively to specific *cis*-elements in *CLE-RS2*, *NF-YB1* and *CRE1* promoters, thereby antagonistically regulating gene expression (Soyano *et al*., 2015; Lin *et al*., 2018; Nishida *et al*., 2021; Luo *et al*., 2022) for example *via* the inhibitory effect of NLP1 on NIN-mediated *CRE1* expression in *M. truncatula* (Lin *et al*., 2018). Likewise in *L. japonicus*, NLP4 accumulates in the nucleus upon nitrate sensing to activate the expression of *CLE-RS2*, which is also activated by NIN upon rhizobial infection (Soyano *et al*., 2014; Nishida *et al*., 2018). The interplay between NIN and NLPs therefore possibly influences not only RNS but also LR formation. Ectopic overexpression of *NIN* potentially disrupts the interaction of the NLPs, thereby interfering with the regulation of LR organ formation.

### Changes in root system architecture as a consequence of multiple independent losses of NIN among species with a loss-of-RNS ancestry

Multiple independent losses or fragmentations of *NIN* and *Rhizobium-directed Polar Growth* (*RPG* (Arrighi *et al*., 2008)) were observed among members of the FaFaCuRo clade (Griesmann *et al*., 2018; van Velzen *et al*., 2018). These losses explain, at least for the species investigated, the inability to engage in the RNS with nitrogen-fixing bacteria. We observed that ectopic expression of *L. japonicus CCaMK^1^*^−*314*^ enhanced the formation of LRs in the RNS-forming species *D. drummondii* to a significantly greater level than in the wild strawberry (*F. vesca*), which has lost *NIN* (Griesmann *et al*., 2018) (Fig. **5**). It is therefore tempting to speculate that *NIN* is also required to mediate this developmental response in non-legume species, for example in *D. drummondii*. The recruitment of the LR developmental program by *NIN* was suggested by the report of NIN-targeted *cis*-regulatory elements within *ASL18* introns; however, these binding sites are only present in legume species (Soyano *et al*., 2019). Our data suggest that the co-option of the LR developmental program by *NIN* extends beyond this family (Figs. **5****, 6**).

### NSP-mediated strigolactone biosynthesis inhibits, while NIN promotes lateral root development

We found that the number of LRs formed by the *nin-2* mutant in our root culture system decreased upon ectopically expressing *CCaMK^1^*^−*314*^ (Fig. **2a**). This finding suggested that CCaMK^1−314^ activates one or multiple repressor(s) of LR formation, which are counteracted by *NIN*. Possible candidates for these repressors are NSP1 and NSP2, which have at least two potential regulatory targets relevant to LR induction. On the one hand, their genes regulate the expression of strigolactone biosynthesis genes such as *Dwarf27* in *M. truncatula*, rice, and barley upon phosphate starvation, leading to a lower LR density at least in rice (Liu *et al*., 2011; Li *et al*., 2022a; Yuan *et al*., 2023). On the other hand, NSP1 and NSP2 were reported to form a complex that interacts with IPN2 and induce *NIN* expression in *Nicotiana benthamiana* leave cells (Xiao *et al*., 2020). It will be interesting to find out which of the two targets plays a role and under which conditions. It is possible that during the CCaMK^1−314^-mediated LR formation, *NIN* regulates the activity of the NSPs which appear to repress the formation of LRs under non-symbiotic conditions (Figs. **4**, **6**). Chiu *et al*. documented a very strong deregulation of LR density by mutation of strigolactone biosynthesis genes or strigolactone receptor genes (Chiu *et al*., 2022). This result is in line with the phenotypes observed in this study for the *nsp1* and *nsp2* mutants, featuring an enhanced LR density (Fig. **4a**).

## Material and Methods

### Plant materials and bacterial and fungal strains

The *Lotus japonicus* ecotype Gifu B-129 was used as the wild type (WT) in this study alongside the mutants *ccamk-3*, *cyclops-3*, *nin-2*, *nsp1-1*, *nsp2-2,* and *snf1-1* (Perry *et al*., 2009). Seed bags are listed in Table **S1**. Seeds from yellow dryas (*Dryas drummondii*, DA462) were purchased from the seed producer Jelitto (Jelitto Staudensamen GmbH, Schwarmstedt, Germany). Seeds from wild strawberry (*Fragaria vesca*) were collected on an excursion to the Breitachklamm in the Kleinwalsertal, Germany.

### *Lotus japonicus* growth conditions and quantification of primary root length and lateral root numbers

*L. japonicus* seeds were scarified, surface-sterilized with 1.2% (w/v) NaClO for 10 min, rinsed five times with autoclaved deionized water, and allowed to soak in water at room temperature for 4 h under agitation. Hydrated seeds were germinated on deionized water solidified with 0.8% (w/v) Bacto agar (Becton Dickinson and Co., Heidelberg, Germany) in square plates (120×120×15 mm). Plates were kept in the dark for 3 days at 24°C before being transferred to low-light conditions in a Panasonic (MLR-352H-PE; **Figs. 1c, 2c, 4a**) growth cabinet at 24°C under a 16-h light/8-h dark photoperiod (50 µmol m^−2^ s^−1^) and grown for 4 more days.

For phenotypic analysis, 7-day-old seedlings were transferred to autoclaved Weck jars (SKU 745, J. WECK GmbH u. Co. KG, Wehr, Germany) containing 300 mL of dry sand:vermiculite mixture (1:1, v/v) and 20 mL of modified Hoagland’s medium (using the “Solution lacking nitrogen” (Hoagland & Arnon, 1950)) and adding 1 mM KNO_3_; 2 mM KH_2_PO_4_; 0.1 µM CoCl_2_; removing Ca(H_2_PO_4_)_2_; replacing H_2_MoO_4_ with 0.1 µM Na_2_MoO_4_; and replacing iron tartrate with 20 µM Na_2_EDTA and 20 µM FeSO_4_; pH 5.8) (for **Figs. 1c, 2c, 4a**). Weck jars were placed in a growth chamber at 24°C under a 16-h light/8-h dark photoperiod (275 µmol m^−2^ s^−1^). Plants were harvested at 30 days post germination (corresponding to 23 days spent in the Weck jars). Harvested plants were examined under a stereomicroscope to score emerged LRs and placed on a flatbed scanner (Epson V700) and imaged at a resolution of 800 dots per inch. Primary root length was measured for each plant with ImageJ (https://fiji.sc).

### *Lotus japonicus* hairy root transformation and hairy root liquid cultures

*L. japonicus* hairy root transformation was performed as described previously (Charpentier *et al*., 2008). The selection of transformed roots was carried out with a green fluorescent protein (GFP) transformation marker encoded on the T-DNA using a Leica M165 FC fluorescent stereomicroscope equipped with a GFP3 filter set (Leica Microsystems, Wetzlar, Germany). Plants with emerging transformed roots were transferred to Fåhraeus medium (Fahraeus, 1957) containing 0.1 μM of the ethylene biosynthesis inhibitor L-α-(2-aminoethoxyvinyl)-glycine and solidified with 1% (w/v) agar (cat. no. HP696, Kalys SA, Bernin, France) in square plates (120×120×15 mm) 2 weeks post inoculation with *Agrobacterium rhizogenes* strain AR1193 (Stougaard *et al*., 1987) and kept on Fåhraeus plates for 2 weeks. Primary root tips of 1.5 cm in length were excised and transferred to round Petri dishes (8.5 cm diameter) containing 18 mL of liquid modified Strullu-Romand (MSR) medium (Declerck *et al*., 1998) (Table **S2**), sealed with Micropore tape (3M Health Care, cat. no. 1530-0) and grown at 22°C in the dark (**Figs. 1–4**; **S2–S5**). The number of emerged LRs (first, second, and higher-order LRs) was scored after 10, 20, or 30 days of incubation (doi) in MSR medium. Sectioning was performed on roots at 10 doi (Fig. **S5**). Roots were embedded in 6% (w/v) low-melting agarose and sliced into 40–50-µm-thick sections using a vibrating-blade microtome (Leica VT1000 S, Wetzlar, Germany). Root sections were imaged with a Leica DM6B fluorescent microscope equipped with a DMC2900 camera (Leica Microsystems, Wetzlar, Germany).

### *Dryas drummondii* and *Fragaria vesca* growth conditions and root transformation

Seeds of *D. drummondii* were germinated as described previously (Billault-Penneteau *et al*., 2019). Seeds of *F. vesca* were transferred to empty mini spin columns (for example GeneJET RNA spin columns, ThermoFisher, Germany) and treated for 5 min with concentrated H_2_SO_4_. Columns were spun in microfuge tubes for 30 sec at 20,000 g to separate the seeds for the H_2_SO_4_, which was removed by pipetting. The seeds were then rinsed five times with sterile deionized water and incubated at 4°C for 5 h under agitation. Seeds were germinated on deionized water solidified with 0.8% (w/v) Bacto agar (Becton Dickinson and Co., Heidelberg, Germany) in square plates (120×120×15 mm) and placed in a Panasonic growth chamber at 22°C under a 16-h light/8-h dark photoperiod (50 µmol m^−2^ s^−1^).

*D. drummondii* hairy root transformation was performed as described previously (Billault-Penneteau *et al*., 2019). Transgenic hairy roots in *F. vesca* were induced by *A. rhizogenes* AR1193. Transformation was performed by cutting 12-day-old *F. vesca* seedlings on filter paper soaked with bacterial suspension. Seedlings were kept in the dark at 18°C for 3 days and then placed in a Panasonic growth chamber at 22°C under a 16-h light/8-h dark photoperiod (50 µmol m^−2^ s^−1^). *A. rhizogenes*-transformed composite *D. drummondii* plants were grown on modified Hoagland’s medium (modified as described above) solidified with 0.4% (w/v) Gelrite (Duchefa, Haarlem, The Netherlands) in square plates (120×120×15 mm). *A. rhizogenes*-transformed composite *F. vesca* plants were grown on Gamborg’s B5 medium (Gamborg *et al*., 1968) solidified with 0.8% (w/v) Bacto agar (Becton Dickinson and Co.) in square plates for 3 days and then transferred to Gamborg’s B5 medium containing 300 µg L^−1^ cefotaxime. Thirty days post *A. rhizogenes* inoculation, *D. drummondii* and *F. vesca* composite plants were screened to identify transformed roots carrying a GFP transformation marker encoded on the T-DNA. For each plant, a single transformed root was kept while all additional untransformed and transformed roots were cut off. The plants were then transferred onto modified Hoagland’s medium (modified as described above) solidified with 0.4% (w/v) Gelrite (Duchefa, Haarlem, The Netherlands). At 45 and 42 days post *A. rhizogenes* inoculation, *D. drummondii* and *F. vesca* composite plants, respectively, were transferred to sterile Weck jars (SKU 745, J. WECK GmbH u. Co. KG, Wehr, Germany) containing 300 g of expanded clay aggregates (2–3 mm diameter; Seramis GmbH, Mogendorf, Germany) and 50 mL of modified Hoagland’s medium (using the “Solution lacking nitrogen” (Hoagland & Arnon, 1950) and adding 100 µM KNO_3_; 250 µM KH_2_PO_4_; 0.1 µM CoCl_2_; removing Ca(H_2_PO_4_)_2_; replacing H_2_MoO_4_ with 0.1 µM Na_2_MoO_4_; and replacing iron tartrate with 20 µM Na_2_EDTA and 20 µM FeSO_4_; pH 5.8) (Fig. **5**). *D. drummondii* and *F. vesca* composite plants were harvested at 20 and 23 days after transfer to the Weck jars, respectively. Roots were scanned and images analysed as described above to assess LR number and primary root length.

### DNA constructs

A detailed description of the constructs and cloning strategies used in this study is provided in Table **S3**. A list of primers can be found in Table **S4**.

### Data visualization and statistical analysis

Statistical analysis and data visualization were performed in R Studio 1.1.383 (RStudio Inc.). Boxplots were used to display data in **Figs. 1–5** and **S3–S4** (black lines, median; box, interquartile range (IQR); whiskers, lowest and highest data point within 1.5 IQR; numbers above or below the whiskers, data points outside the 1.5 × IQR). The R package “ggplot2” (Wickham, 2009) was used to generate the boxplots. Statistical results are displayed in lowercase letters where different letters indicate statistical significance. Tests applied are stated in the figure legends.

## Acknowledgements

We thank Rosa Elena Andrade for providing the RAp15 plasmid (Table **S3**). MP acknowledges funding for this project from the European Research Council (ERC) under the European Union’s Seventh Framework Programme (FP7/2007-2013) under grant agreement n° 340904 (EvolvingNodules), from the Deutsche Forschungsgemeinschaft (DFG) - Projektnummer 170483403, in the context of the SFB924 ‘Molecular basis of agronomic traits’, and from the ANR-DFG project ‘COME-IN’ Deutsche Forschungsgemeinschaft (DFG) - Projektnummer 258665719.

## Author contributions

CC generated hairy root liquid cultures and root phenotyping presented in **Figs. 1, 2, 3, 4a** and **c**, **5, S3**, **S4** and **S5b**. AS performed hairy root transformation in *D. drummondii* and *F. vesca* presented in Fig. **5**. MKR-L modified the MSR medium, established the experimental setup for hairy root liquid cultures, and acquired preliminary data and data presented in Figs. **4b****, S1** and **S5a**. CC, MKR-L, and MP conceived the research, designed the experiments, and analyzed the data. MKR-L and CC prepared the figures. CC, MKR-L and MP wrote the manuscript.

## Declarations of interests

The authors declare no competing interests.

## Notes

### Competing Interest Statement

The authors have declared no competing interest.

