## SupplementaryFigures for "Lateral root formation is stimulated by common symbiosis genes and *NIN* in *Lotus japonicus*"

**Supplementary Figures**

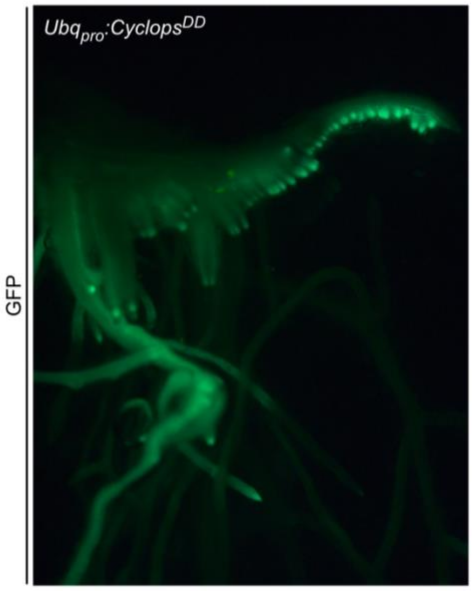


**Figure S1: Ectopic and constitutive expression of *Cyclops*$^{DD}$ increases lateral root number in transgenic roots of composite *Lotus japonicus* plants.** Picture of a transgenic root transformed with *Ubq*$_{pro}$:*Cyclops*$^{DD}$ from a composite *L. japonicus* WT plant. The transgenic root expressed free GFP as a visual transformation marker. Note the densely packed emerging lateral organs.

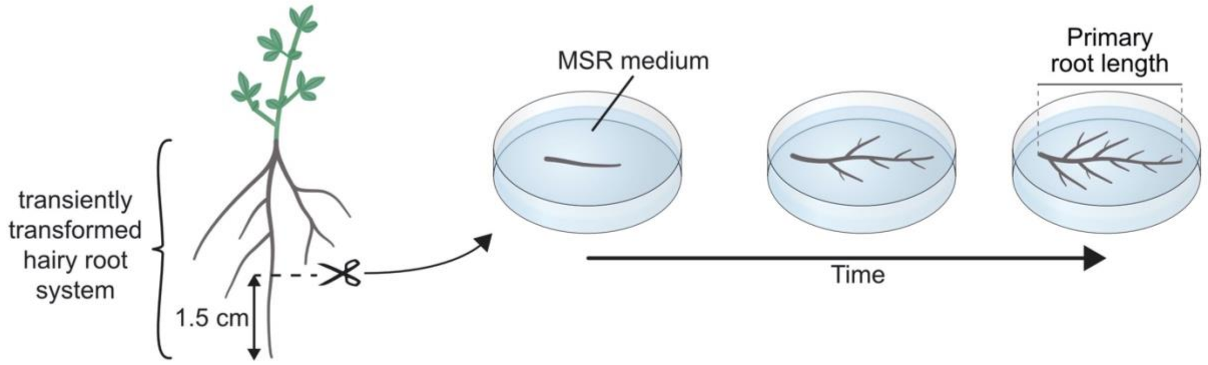


**Figure S2: Diagram of the experimental setup for cultivating hairy root liquid cultures.** Primary root tips (1.5 cm) of 4 weeks old transformed *L. japonicus* hairy roots were cut off and transferred to Petri dishes containing 18 mL of liquid modified Strullu-Romand (MSR) medium. Plates were sealed with micropore tape and placed at 22 °C in the dark. Emerged lateral roots were counted after 10, 20 or 30 days of incubation in the MSR medium.

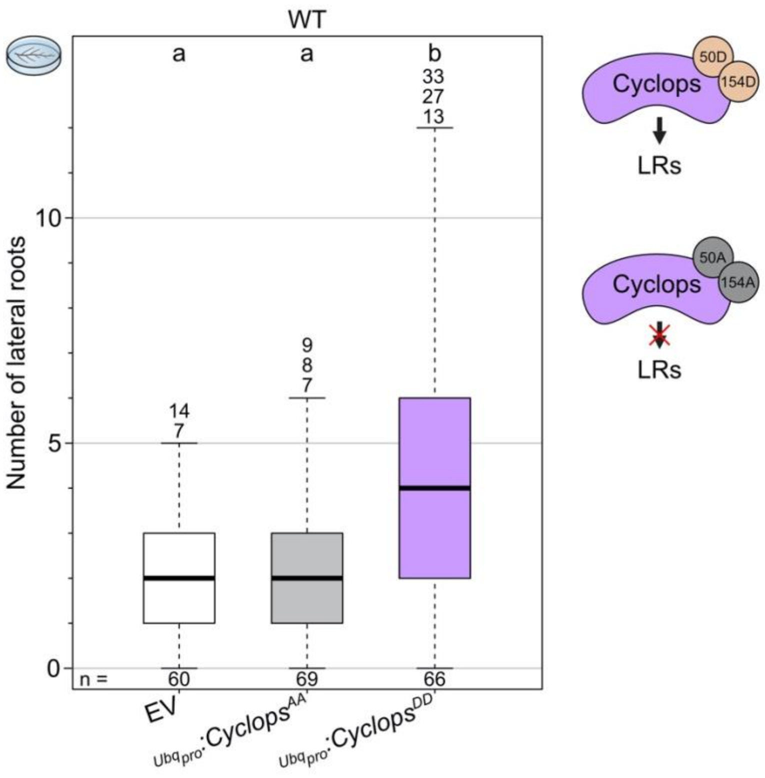


**Figure S3. Expression of a phosphomimic version of Cyclops, but not of its phosphoablative version, stimulates lateral root formation.** Hairy root liquid cultures of *L. japonicus* WT roots transformed with the empty vector (EV), *Ubq$_{pro}$:Cyclops$^{AA}$* or *Ubq$_{pro}$:Cyclops$^{DD}$*. Box plots represent the number of lateral roots per root culture after 10 days of incubation. Ectopic expression of phosphoablative Cyclops$^{AA}$ does not increase lateral root numbers. Data were subjected to a Kruskal-Wallis test followed by Dunn's *post hoc* analysis; $p < 0.05$. n: number of roots analyzed.

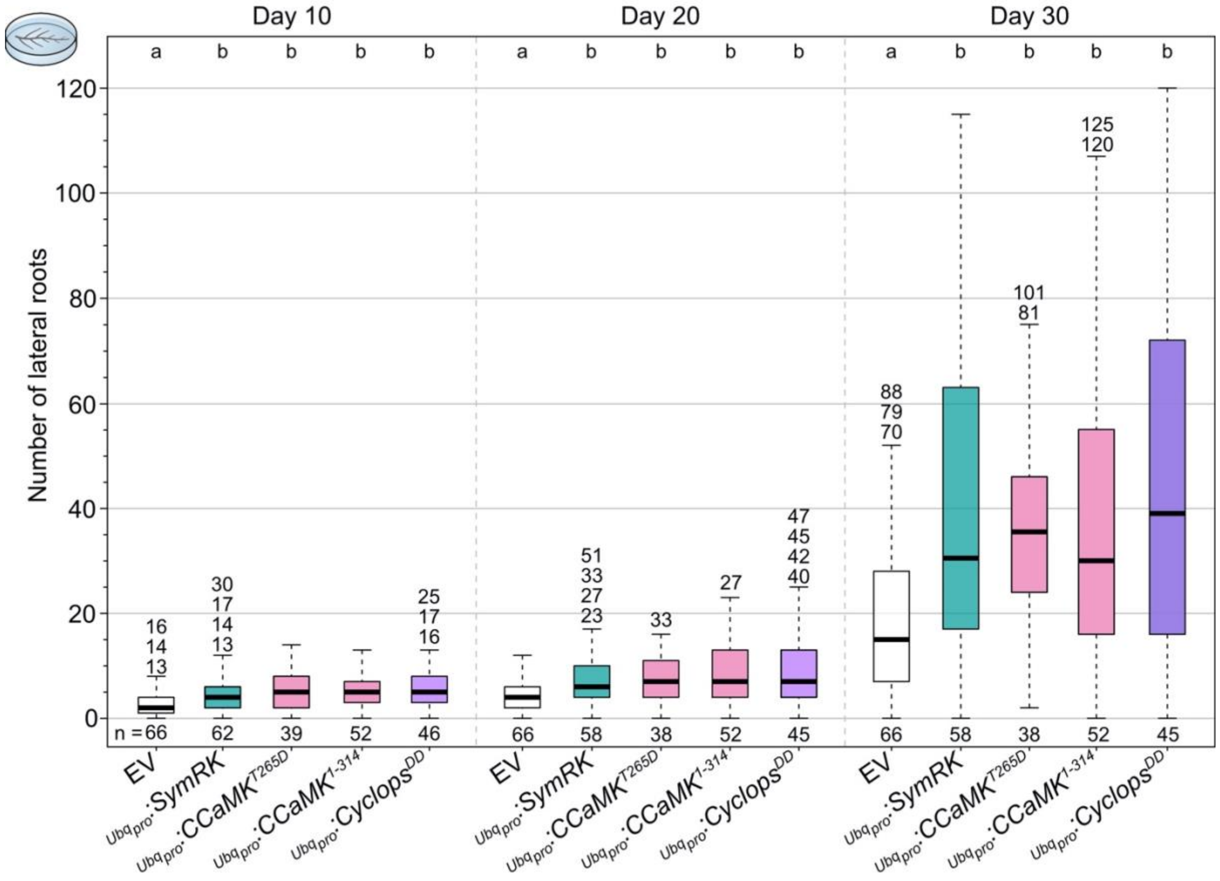


**Figure S4. Time-course analysis of lateral root numbers in root cultures expressing deregulated versions of CCaMK or Cyclops or ectopically overexpressing *SymRK*, (related to Fig. 1).** Hairy root liquid cultures of *L. japonicus* WT roots transformed with the empty vector (EV), *Ubq$_{pro}$:SymRK*, *Ubq$_{pro}$:CCaMK$^{T265D}$*, *Ubq$_{pro}$:CCaMK$^{1\text{-}314}$* or *Ubq$_{pro}$:Cyclops$^{DD}$*. Box plots represent the number of lateral roots per root culture after 10, 20 and 30 days of incubation. Note that root cultures expressing deregulated versions of *CCaMK* or *Cyclops* or ectopically overexpressing *SymRK*, have more lateral roots than those transformed with the EV at all analyzed time points. Data were subjected to a Kruskal-Wallis test followed by Dunn's *post hoc* analysis; $p < 0.05$. n: number of roots analyzed.

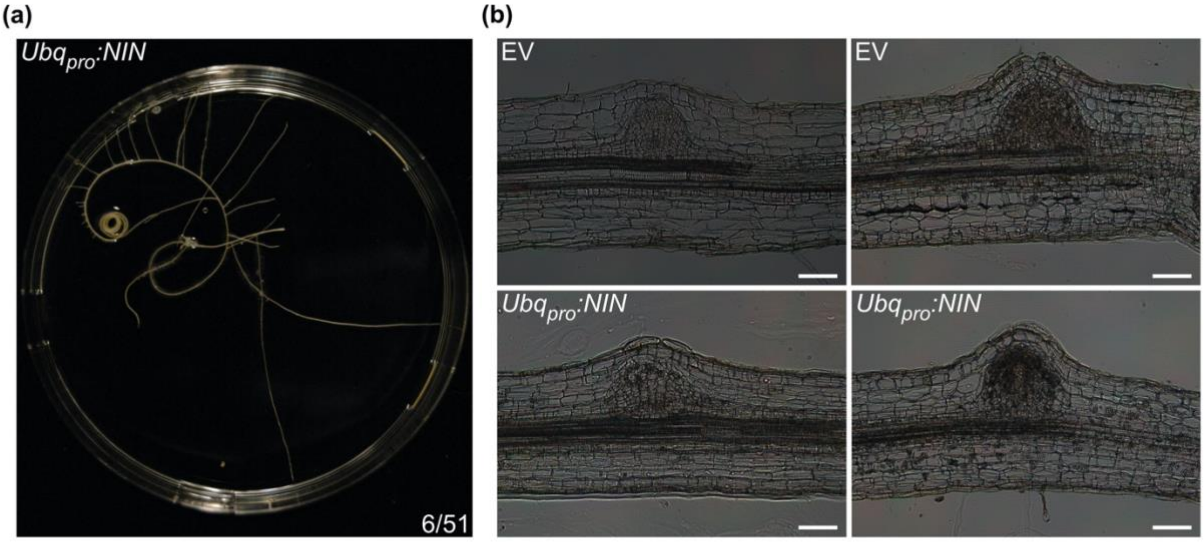


**Figure S5. Ectopic expression of *NIN* in hairy root liquid culture results in excessive curling (related to Fig. 2).** (**a**) Representative picture of an excessively curling *L. japonicus* WT root transformed with *$Ubq_{pro}$:NIN* after 20 days of incubation in MSR liquid medium. Numbers: roots exhibiting excessive curling/total roots analyzed. (**b**) Pictures of root sections of *L. japonicus* WT transformed with the empty vector (EV) or with *$Ubq_{pro}$:NIN* after 10 days of incubation. Note that lateral roots formed on roots ectopically overexpressing *NIN* were anatomically similar to those observed on roots transformed with the EV. This phenomenon was also reported in *M. truncatula* roots ectopically expressing *LBD16* (Schiessl *et al.*, 2019). Bars, 100 µm.
